# Hypoxia Inducible Factor 1α-driven steroidogenesis impacts systemic hematopoiesis

**DOI:** 10.1101/2025.03.20.644292

**Authors:** Deepika Watts, Nicolas Eberz, Mangesh T. Jaykar, Anupam Sinha, Cagdas Ermis, Johanna Tiebel, Ulrike Baschant, Martina Rauner, Tatyana Grinenko, Triantafyllos Chavakis, Peter Mirtschink, Ali El-Armouche, Ben Wielockx

## Abstract

Glucocorticoids regulate hematopoiesis, but how chronic elevation of endogenous glucocorticoid production affects hematopoietic stem cell (HSC) function and immune cell development remains incompletely understood. Using an adrenocortical cell-specific HIF1α (Hypoxia inducible Factor-1α)-deficient mouse model (P2H1^Ad.Cortex^) resulting in elevated glucocorticoid (GC) levels, we here demonstrate that sustained GC exposure promotes hematopoietic stem and progenitor cell (HSPC) expansion while shifting HSCs toward a more quiescent and metabolically restrained state. Functionally, these HSCs exhibited enhanced regenerative potential, as evidenced by superior donor chimerism in transplantation assays. In addition, we observed a striking increase in myeloid progenitors, as well as in their progeny (monocytes and granulocytes). Conversely, B-cell differentiation in the bone marrow was severely impaired, with a strong block at the pre-pro-B cell stage.

To determine whether these phenotypes were driven by glucocorticoid receptor (GR) signaling, we performed transplantation experiments using GR-deficient or WT control bone marrow into P2H1^Ad.Cortex^ or WT littermate recipients. This approach decisively demonstrated that both the increase in myeloid cells and the block in B-cell differentiation were GR-dependent, confirming that GC-GR signaling plays a pivotal role in shaping hematopoiesis.

Taken together, our findings clearly suggest a direct role for chronic glucocorticoid exposure in regulating HSC function, lineage differentiation, and stress hematopoiesis. The mouse model of adrenocortical cell-specific HIF1α deficiency provides a valuable tool to study the long-term effects of elevated glucocorticoid levels on hematopoietic regulation and may provide further insight into hematologic disorders associated with chronic therapeutic glucocorticoid administration.

**Article Summary:**

– Our study aimed to elucidate how chronic elevation of glucocorticoids impacts hematopoietic stem cell function and immune cell development using an adrenocortical cell-specific HIF1α-deficient mouse model.
– The main conclusion is that sustained glucocorticoid exposure, through glucocorticoid receptor signaling, promotes hematopoietic stem and progenitor cell expansion with enhanced regenerative potential while skewing lineage differentiation toward myeloid expansion and impeding B-cell development.

## Introduction

Hematopoiesis, the process responsible for generating blood cells, is fundamental to the biological system of all vertebrates. It depends primarily on hematopoietic stem cells (HSCs), a rare and specialized population of cells in the bone marrow, where they maintain lifelong blood production through self-renewal and differentiation into all blood and immune lineages.^1, 2^ HSCs are rare and reside in niches where oxygen levels are significantly lower than in other tissues.^3, 4^ Beyond hypoxic conditions, the bone marrow microenvironment, which includes endothelial cells, perivascular cells, and mesenchymal stem cells, plays a crucial role in regulating HSC maintenance and function.^5, 6^ Through continuous interactions with these niche components and systemic signals, hematopoiesis remains highly dynamic, adapting to physiological demands and external stimuli.^7^ External factors such as stress, inflammation, infection, and aging further modulate this process, often driving stress-induced hematopoiesis.^7–10^ However, the long-term effects of chronic physiological stress on hematopoietic function remain incompletely understood.

The adrenal gland plays a critical role in stress adaptation by regulating steroidogenesis, which affects immune function and systemic homeostasis.^11, 12^ Glucocorticoids (GCs) are widely used in immunosuppressive therapies, particularly in inflammatory and autoimmune diseases.^13^ Our previous findings showed that modulation of HIF1α expression/activity in the adrenal cortex dramatically alters adrenal steroidogenesis, leading to systemic changes in circulating glucocorticoids and inflammatory cytokines.^14^ This raises the important question of how chronic increases in endogenous glucocorticoid production affect HSC and immune cell development, especially when compared to the well-documented effects of synthetic glucocorticoids such as dexamethasone, when administered therapeutically.^15–18^

Furthermore, with the increasing interest in HIF stabilizers as potential therapeutic agents, understanding their effects on adrenal steroidogenesis and hematopoiesis is crucial.^19^ Our mouse model with HIF1α deficiency in adrenocortical cells (Akr1b7:cre-*PHD2/HIF1*^ff/ff^ (P2H1^Ad.cortex^))^14^, accompanied by higher glucocorticoid levels, provides a unique tool to mimic chronic stress, allowing us to evaluate the effects of elevated endogenous glucocorticoid levels on hematopoietic function. This model also provides insights into the potential adverse effects of HIF inhibitors on adrenal steroidogenesis and immune regulation.

In this study, we show that HIF1α deficiency in the adrenal cortex leads to systemic glucocorticoid overproduction, which promotes HSPC expansion while enforcing a more quiescent HSC state at steady state. These HSCs exhibit a competitive advantage in repopulating the bone marrow after irradiation. In addition, we reveal the complex effects of chronic enhanced glucocorticoids on myeloid and lymphoid maturation. Our findings highlight the indispensable role of HIF1α in systemic physiological regulation and position our mouse model as a valuable tool for understanding the pathophysiological consequences of glucocorticoid dysregulation in human health and disease.

## Methods

### Mice

Experiments were performed in male and female mice up to one year of age. Akr1b7:cre-*Phd2*/*Hif1^ff/ff^*(P2H1^Ad.Cortex^ or P2H1) and Akr1b7:cre-*Phd2*/*Phd3^ff/ff^*(P2P3^Ad.Cortex^ or P2P3) mice were generated as previously reported by us.^14^ C57BL/6 (B6) and B6.SJL-PtprcaPep3b/BoyJ (SJL) mice were crossed to generate F1 progeny (CD45.1/CD45.2) for transplantation experiments. B6.Cg-Nr3c1tm1.1Jda/J (GR^f/f^; C57BL/6J) mice were purchased from The Jackson Laboratory^20^ and crossed to B6.Cg-Tg(VAV1-cre)1Graf/MdfJ (VavCre; C57BL/6J) mice^21^ to produce Vav:cre-*GR^f/f^* (GR^HSC^) mice. All mice were bred as homozygous for GR^f/f^ and heterozygous for Cre. Cre-negative GR^f/f^ mice were used as WT littermates to GR^HSC^. All mice were maintained under specific pathogen-free (SPF) conditions. All mice were born in a normal Mendelian manner. For each experiment, transgenic mice were compared with littermate controls (Cre-negative). Mice were genotyped using described primers.^14^ Both genders of mice were used in comparable numbers, and no significant differences were observed between genders of the same genotype for any of the analyses performed within this study. All mice were bred and maintained in accordance with animal welfare guidelines and protocols approved by the Landesdirektion Sachsen, Germany.

### Blood analysis

Peripheral blood was drawn from mice by retro-orbital sinus puncture using heparinized micro hematocrit capillaries (VWR, Darmstadt, Germany) and blood was further diluted with PBS (1:5) according to the user manual. White blood cells, red blood cells and platelets were measured in whole blood using a Sysmex automated blood cell counter (Sysmex 117 XE-5000), and plasma was separated and stored at −80 °C.

### Flow cytometry and single cell sorting

Single-cell suspensions were prepared from bone/bone marrow, lymph nodes, thymus and spleen samples by mechanical disruption and filtration through 70 μm cell strainers (Becton Dickinson, San Diego, CA, USA). The cell suspension from bone marrow and spleen samples was subjected to red blood cell lysis with ACK lysis buffer (Life Technologies Cat. A10492-01). The single-cell suspensions were then stained with the appropriate antibodies. Prior to staining, the cells were incubated with the bio mix of biotinylated antibodies for the BM and in the next round stained with fluorophore-conjugated antibodies. All antibodies were either from ThermoFischer Scientific (eBioscience); BioLegend or BD Biosciences (Supplementary Table I). Cell sorting was performed using BD Aria II or III as described previously.^22^

### Apoptosis and Cell cycle

For apoptosis analysis, cells were first surface-stained, washed twice with 200 µl cold PBS, and then resuspended in 1ml 1X Binding Buffer. Annexin V (556420) (BD-Pharmingen) was then added to 100ul of suspension and cells were incubated for 20 minutes in the dark at room temperature, followed by DAPI staining. For analysis, cells that were negative for DAPI and Annexin V were considered to be alive, while cells that were positive for DAPI and Annexin V were considered to be dead. Furthermore, Annexin V-positive, DAPI-negative cells were considered to be undergoing apoptosis. Cell Cycle analysis was performed as described previously. ^23^ Briefly, HSCs were fixed, permeabilized and stained for intracellular Ki-67 (PE, B56, BD Biosciences) to differentiate between G0 and G1 phases. DAPI was used to measure DNA content and to separate cells in S/G2/M phase from G0 and G1 cells.

### Transplantation

For HSC transplantation, HSCs (cKit^+^Sca-1^+^CD48^-^CD150^+^) were isolated from the BM of P2H1^Ad.Cortex^ and WT littermates using BD fluorescence-activated cell sorting (Aria II). Cells were competitively transplanted together with 5 x 10^5^ B6.SJL (CD45.1) total BM competitor cells into lethally irradiated (9 Gy) F1 recipients. For the GC-GR transplantation experiments, lethally irradiated recipient P2H1^Ad.Cortex^ and WT littermates were iv injected with 5×10^6^ Erythrolyzed total BM cells from GR^HSC^ or WT. BM, spleen or peripheral blood we analyzed at different time points as indicated in the text using BD LSR Fortessa™ Cell Analyzer or BD FACSymphony-A5.

### Transcriptome Mapping

Low quality nucleotides were removed using Illumina fastq filter (http://cancan.cshl.edu/labmembers/gordon/fastq_illumina_filter/). Reads were further subjected to adaptor trimming using cutadapt.^24^ Alignment of the reads to the Mouse genome was done using STAR Aligner^25^ using the parameters: “--runMode alignReads --outSAMstrandField intronMotif --outSAMtype BAM SortedByCoordinate --readFilesCommand zcat”. Mouse Genome version GRCm38 (release M12 GENCODE) was used for the alignment.

### Read Quantification

Using the parameters: ‘htseq-count -f bam -s reverse -m union -a 20’, HTSeq-0.6.1p1^26^ was used to count the reads that map to the genes in the aligned sample files. The GTF file (gencode.vM12.annotation.gtf) used for read quantification was downloaded from Gencode (https://www.gencodegenes.org/mouse/release_M12.html).

### Differential Expression Analysis

Gene centric differential expression analysis was performed using DESeq2_1.8.1.^27^ Also, the raw read counts for the genes across the samples were normalized using ‘rlog’ command of DESeq2 and subsequently these values were used to render a PCA plot using ggplot2_1.0.1.^28^ Heatmaps were generated using Complex Heatmap package of R/Bioconductor.^29^

### Functional Analyses

Pathway and functional analyses was performed using GSEA^30^ and EGSEA.^31^ GSEA/EGSEA were run using normalized gene expression matrix against databases like Molecular Signatures Database (MSigDB), Reactome, KEGG and GO based repositories.

### Pathway Activity

Additionally, the pathway activity calculations were performed using progeny^32^. A R package progeny is a compendium of 14 pathways, which have been extracted from publicly available perturbation experiments. The input to the progeny() command was the normalized gene expression matrix.

### Statistical analysis

To assess statistical significance between two experimental groups, a Mann–Whitney U-test was used for non-normally distributed data, while an unpaired t-test with Welch’s correction, accounting for unequal variances, was applied for normally distributed data. Normality was tested using the Shapiro-Wilk test. Statistical differences presented in the figures were considered significant at p-values below 0.05. All statistical analyses were performed using GraphPad Prism v10.02 for Windows or higher (GraphPad Software, La Jolla, California, USA, www.graphpad.com).

## Results

### HIF1α-dependent chronic glucocorticoid exposure promotes HSPC expansion and enforces HSC quiescence via p53 pathway activation

Previously, we demonstrated that chronic inhibition of HIF1α in the adrenal cortex enhances steroidogenesis within the gland, leading to increased systemic steroid levels, including glucocorticoids.^14^ Recognizing the pivotal roles of glucocorticoids in immune regulation, stress response, and hematopoiesis, we aimed to investigate the chronic effects of HIF1α-dependent increased steroidogenesis on the hematopoietic system. To this end, we analyzed the bone marrow hematopoietic stem and progenitor cell (HSPC) populations in our P2H1^Ad.Cortex^ mice using flow cytometry (FACS) and a well-established panel of markers (Figure 1A). Interestingly, our analyses reveal a slight but significant increase frequency of HSCs (Lin Kit Sca-1 CD48 CD150) and multipotent progenitors (MPPs) in P2H1^Ad.Cortex^ mice compared to WT littermate controls. Specifically, we observed an increase in MPP2 (Lin Kit Sca-1 CD48 CD150) and MPP3/4 (Lin Kit Sca CD48 CD150), both as a percentage of single cells (Figure 1B, C) and in total cell numbers (Figure S1A, B). To further explore the behavior of these HSCs, we performed bulk RNA sequencing on sorted HSCs from both WT and P2H1^Ad.Cortex^ mice (Figure S1C). Gene set enrichment analysis (GSEA) revealed a pronounced decrease in cell cycle activity and a dramatic reduction in metabolism in P2H1^Ad.Cortex^ HSCs, with no observed changes in cell survival/apoptosis (Figure 1D and Figure S1D). Consistent with these findings, we detected a significant reduction in SCF ubiquitin ligase complex activity, suggesting reduced degradation of key cell cycle inhibitors p27 and p21. In addition, we found a dramatic reduction in RUNX1-associated HSC differentiation, which strongly suggests increased HSC quiescence (Figure 1E). This coupled with increased HSC numbers indicates that these cells, while chronically less proliferative, exhibit a preference for self-renewal over differentiation. Interestingly, we also performed cell cycle analysis using FACS but observed no significant differences in the proportions of HSCs in G_0_ or non-G_0_ phases between WT and P2H1^Ad.Cortex^ HSCs (Figure S1E). Although this snapshot of cell cycle status may seem at odds with the RNAseq data, it is very well possible that FACS does not capture the cumulative effects of chronic steroid exposure, which are better reflected in the RNAseq results and the observed increase in HSC numbers in P2H1^Ad.Cortex^ mice. In support of all observations that reflect the chronicity of our phenotype, a carnival analysis of the RNA sequencing data highlighted the strong involvement of the p53 pathway, which is known to maintain genomic integrity and promote cell stability by preventing excessive proliferation (Figure 1F).^33^ Indeed, within this pathway we identified the modulation of a number of different genes associated with a more quiescent stage of the HSCs. For example, the upregulation of *Trp53bp2* and *Slamf1 (CD150)* supports the quiescent state of HSCs. Trp53bp2 enhances p53-mediated cell cycle arrest and genomic stability, while Slamf1 (CD150) reinforces dormancy by mediating niche interactions and suppressing activation signals. In parallel, the downregulation of *Hsd11b1, Mastl, E2f1* and *Cdk5rap1* contributes to the same outcome, collectively forming a regulatory network that maintains HSC quiescence under chronic glucocorticoid exposure. The suppression of cell cycle regulators Mastl and E2f1 increases quiescence by delaying mitotic entry and inhibiting the G1/S transition. Downregulation of Cdk5rap1 reduces mitochondrial activity, contributing to a low-energy state, while Hsd11b1 downregulation, likely due to chronically elevated systemic glucocorticoid levels, may limit intracellular GC activation and prevent HSC hyperactivation (Figure 1G). Taken together, these combined changes may provide a regulatory network to maintain HSC quiescence under chronic glucocorticoid exposure in P2H1^Ad.Cortex^ mice compared to their WT littermates.

**Figure 1.**
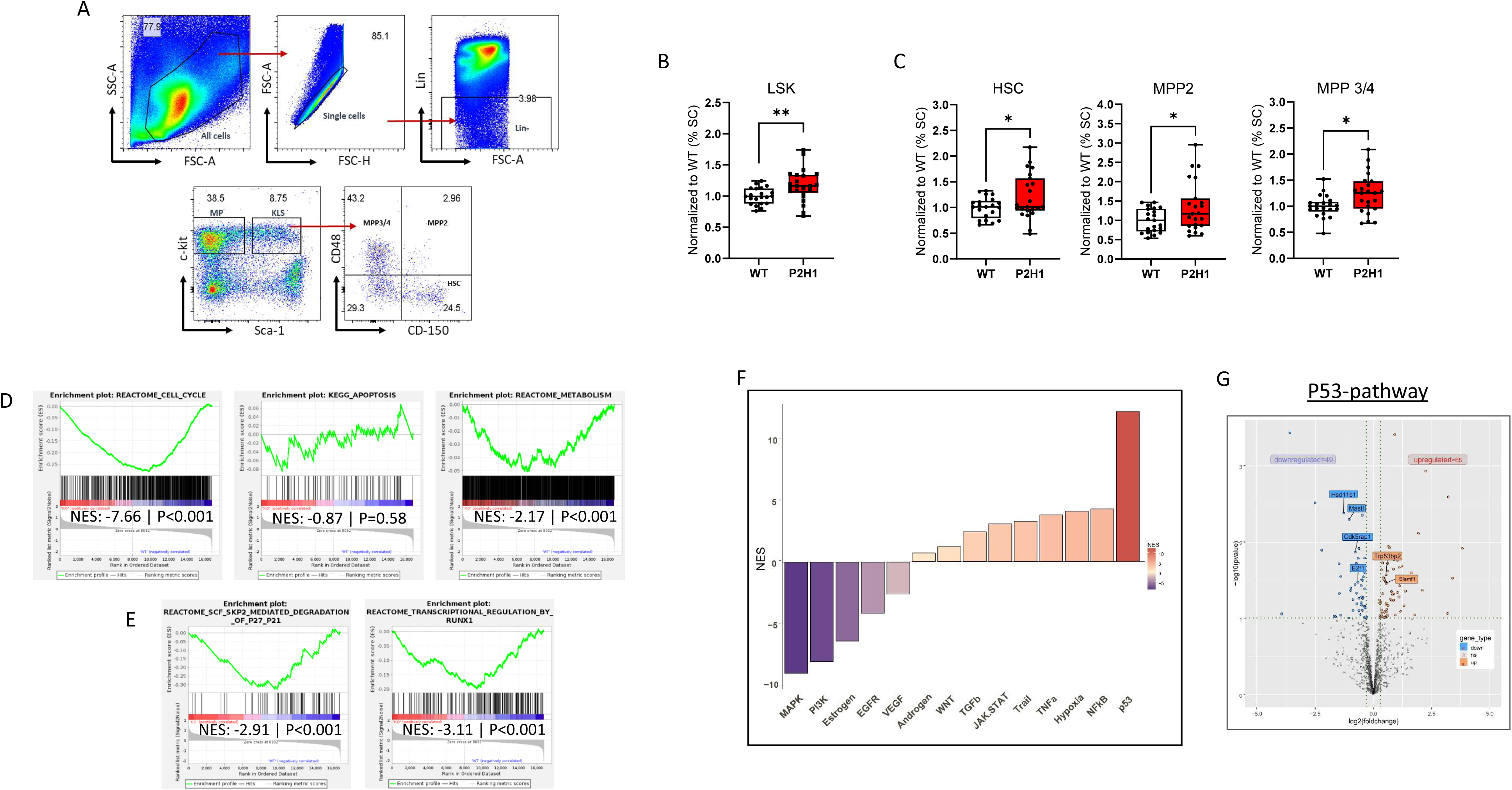
HIF1α-associated steroidogenesis affects hematopoietic stem cell dynamics. (A) FACS gating strategy for identification of HSPC by staining surface markers as detailed in the Methods section. (B-C) Normalized percentage of single cells in the BM of WT mice and Akr1b7:cre-*PHD2/HIF1*^ff/ff^ (P2H1) littermates. LSK (Linage^-^Sca-1^+^cKit^+^) represents HSPCs. Data points represent individual mice from at least three different experiments - normalized values against WT control. Data are represented as box & whisker plots showing all data points with whiskers from min to max. Statistical significance was determined using a Mann–Whitney U-test or unpaired t-test with Welch’s correction (*p<0.05; **p<0.005). (D) GSEA shows significantly decreased cell cycle and metabolism gene signatures and (E) differential changes in the regulation of central cell cycle regulators. (F) Carnival analysis displaying pathway activity scores as Normalized Enrichment Score (NES). (G) A volcano plot depicting the genes associated with the p53-pathway including a selection of up/down regulated genes.

### HSC transplantation reveals enhanced regenerative potential of quiescent P2H1^Ad.Cortex^ HSCs

To validate these findings, we analyzed the remaining HSC function from both genotypes by transplanting these stem cells into irradiated recipient mice to assess their functional capacity to repopulate the bone marrow. We transplanted 300 donor HSCs (CD45.2) together with 5 × 10 competitive bone marrow (BM) cells (CD45.1) into irradiated WT (F1) recipient mice (CD45.1 & CD45.2). Donor-derived white blood cell (WBC) chimerism was then monitored monthly starting four months post-transplantation (Figure 2A). Notably, P2H1^Ad.Cortex^-derived HSCs showed a marked increase in chimerism, particularly in circulating PMNs and T lymphocytes, compared to WT controls (Figure 2B). At six months post-transplant, bone marrow analysis of recipient mice revealed a significant increase in P2H1^Ad.Cortex^-derived CD45 mature cells, consistent with the changes observed in the circulation (Figure 2C). In addition, analysis of HSPC fractions in the BM revealed higher levels of MPP3/4 populations, with a slight decrease in both HSCs and MPP2 populations (Figure 2D). Taken together, these results support our RNA sequencing data and suggest that the more quiescent P2H1^Ad.Cortex^ HSCs experience less exhaustion and, as a result, have a greater regenerative potential to replenish irradiated bone marrow compared to WT HSCs.

**Figure 2.**
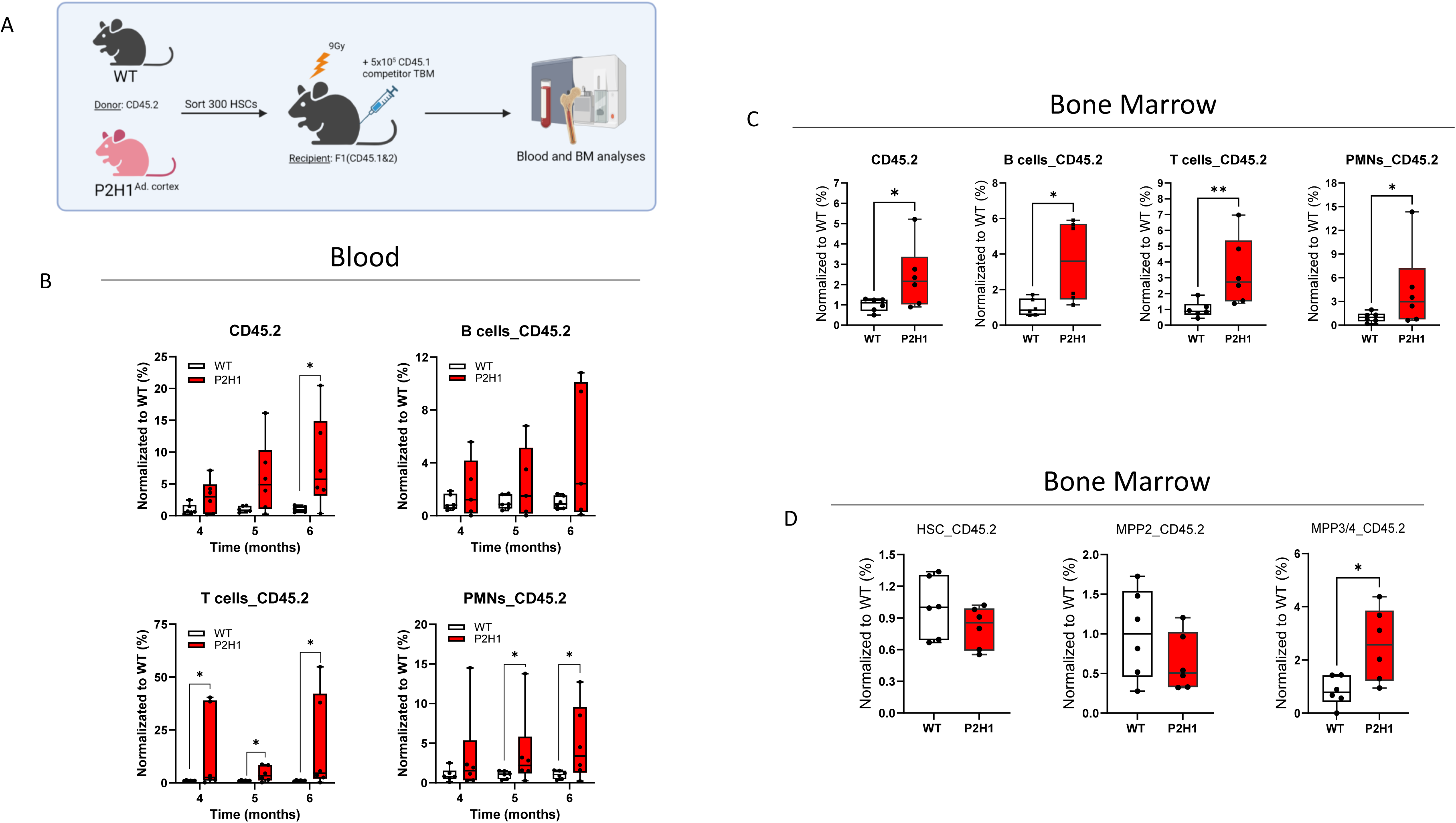
Enhanced replenishment of the hematopoietic system with transplanted hematopoietic stem cells from P2H1 mice. (A) Schematic overview of the HSC-transplantation in irradiated recipients and subsequent blood analyses from 4 months after transplantation and an additional BM analysis at 6 months. (B) Blood analysis and (C-D) BM FACS analysis of mature hematopoietic populations and HSPCs. Data points represent individual mice from at least two different experiments - normalized values against the average of WT controls. Data are presented as box & whisker plots showing all data points with whiskers from min to max. Statistical significance was determined using a Mann–Whitney U-test or unpaired t-test with Welch’s correction (*p<0.05; **p<0.005).

### Increased erythropoiesis independent of bone marrow hematopoiesis

Recent studies have shown that chronic exposure to excess glucocorticoids can lead to overproduction of red blood cells (RBCs) in patients with Cushing’s disease.^34^ Supported by these findings, we examined blood cells in our P2H1^Ad.Cortex^ mouse model and found similar changes. Specifically, our data show that mice chronically exposed to glucocorticoids exhibit a significant increase in RBC numbers, hemoglobin levels and their mean corpuscular volume (MCV) (Figure 3A). We further investigated whether this increase was due to a biased shift towards erythroid progenitors. Analyses of these fractions in the BM (Figure 3B) revealed no significant differences in the populations of bipotent pre-megakaryocyte erythroid progenitors (Pre-MgE), upstream of the more committed erythroid-restricted progenitors (Pre-CFUe) and erythroid colony-forming units (CFUe) (Figure 3C). Similarly, erythroblasts (EBs), the immediate precursors of RBCs in the bone marrow, showed no significant variation in P2H1^Ad.Cortex^ mice compared to WT littermate controls (Figure 3C). Conversely, and consistent with findings from another erythrocytotic mouse model previously studied by our team,^35^ it is the spleen of P2H1^Ad.Cortex^ mice that engages in stress erythropoiesis, containing significantly more erythroblasts than their WT counterparts (Figure 3D). In contrast to the previous study in which erythrocytosis was directly induced by increased erythropoietin (EPO) production,^35^ our P2H1^Ad.Cortex^ mice exhibited significantly lower EPO levels in circulation. This decrease might underlie a negative feedback mechanism possibly induced by the increased number of circulating erythrocytes and the greater potential to deliver oxygen to EPO-producing cells in the kidney, automatically leading to lower HIF2α activity and consequent *Epo* transcription (Figure 3E). These findings strongly suggest that the spleen engages in a mild stress erythropoiesis following long-term exposure to chronically elevated glucocorticoids and corroborate the increased number of splenic erythroblasts observed in P2H1^Ad.Cortex^ mice.

**Figure 3.**
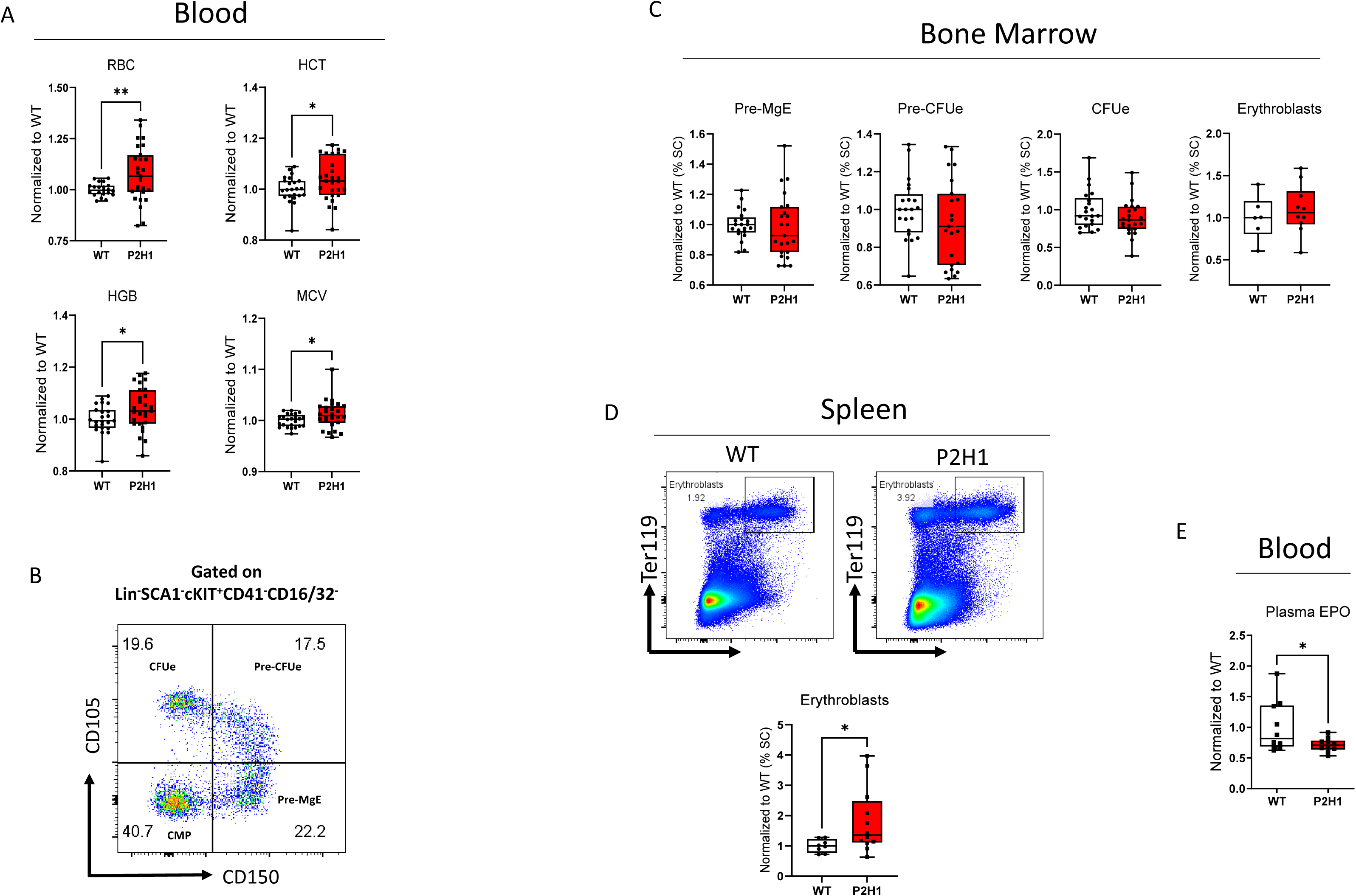
Chronic exposure to glucocorticoids modulates erythropoiesis in P2H1 mice. (A) Normalized number of different RBC parameters in circulation from WT mice and P2H1 littermates under steady state. (B) Representative FACS gating strategy for the identification of erythroid progenitors by staining surface markers as detailed in the Methods section. (C) Normalized percentage of single cells representing different erythroid progenitors in the BM. (D) Representative FACS gating strategy of erythroblasts in the spleen and significantly increased spleen EB single cell percentages in P2H1 mice. (E) EPO ELISA with plasma samples from WT and P2H1 littermates demonstrating significant decrease of EPO levels in circulation. Data points represent individual mice from at least two different experiments - normalized values against the average of WT controls. Data are presented as box & whisker plots showing all data points with whiskers from min to max. Statistical significance was determined using a Mann– Whitney U-test or unpaired t-test with Welch’s correction (*p<0.05; **p<0.005).

### HIF mediated chronic increase in steroidogenesis results in modulation of myelopoiesis

Next, we investigated the effects of prolonged chronic exposure to elevated glucocorticoids on myelopoiesis. First, we analyzed myeloid precursors in the bone marrow and found a significant reduction in common myeloid progenitors (CMPs), alongside a notable increase in granulocyte-monocyte progenitors (GMPs) (Figure S2A, Figure 3B and 4A). This increased GMP levels correlated with a slight but significant increase in the number of monocytes and granulocytes, although eosinophils in this compartment were markedly reduced (Figure S2B-C and Figure 4B). Correspondingly, we observed an increase in circulating monocytes and a decrease in splenic eosinophils (Figure S2D, E). The latter is consistent with previous findings that glucocorticoids can directly regulate eosinophil functionality and survival, potentially leading to lower eosinophil counts.^36^

**Figure 4.**
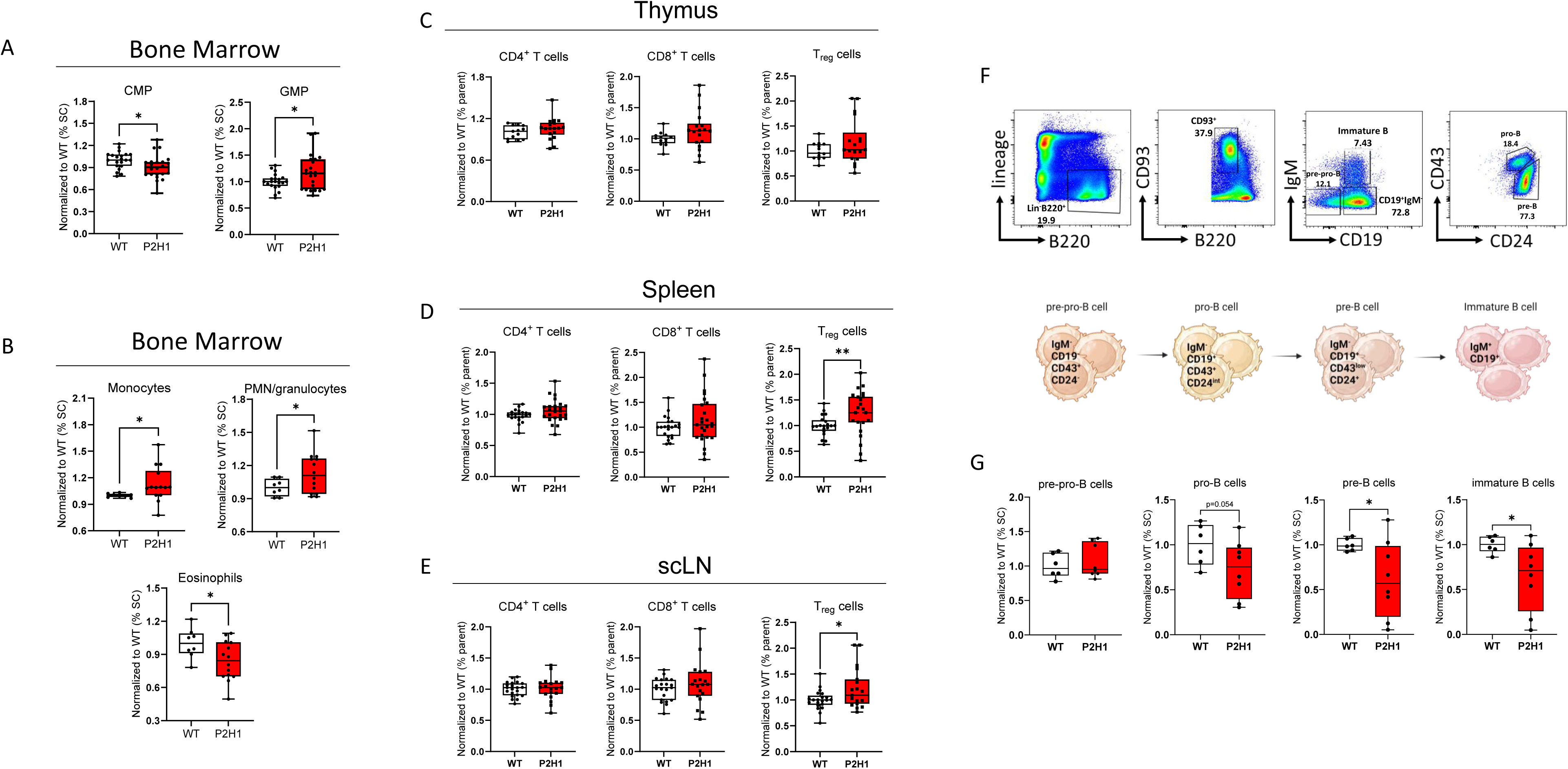
Chronic increased steroidogenesis results in alterations in myeloid cells and lymphocytes. (A-B) FACS analysis of BM-derived myeloid progenitors and mature populations (Normalized from the % of single cells). (C-E) FACS analysis of different T-cell populations in distinct organs (Normalized from the % of parent). Each graph represents data from at least three independent experiments. (F) Representative FACS plots of four different B-cell maturation stages and (G) FACS analysis of BM-derived B-cell progenitors (Normalized from the % of single cells). Data points represent individual mice from at least two different experiments - normalized values against WT control. Data are presented as box & whisker plots showing all data points with whiskers from min to max. Statistical significance was determined using a Mann–Whitney U-test or unpaired t-test with Welch’s correction (*p<0.05; **p<0.005).

### Increased Treg cells in the peripheral lymphoid organs

Previous studies have demonstrated that glucocorticoids significantly affect the function and proliferation of regulatory T cells (Tregs), highlighting their importance in the management and treatment of various immune-related diseases.^37^ Building on this, we extended our study to assess the impact of HIF1α-mediated chronic glucocorticoid elevation by analyzing different T cell lineages in both the thymus and peripheral organs, with a specific focus on CD4+ and CD8+ T cells, and Tregs, identified as CD4^+^CD25^+^Foxp3^+^, in particular (Figure S2F). Interestingly, while no differences in CD4^+^ or CD8^+^ T cell fractions were observed between the organs we examined, Tregs were significantly increased in the spleen and the subcutaneous lymph nodes (scLN) (Figure 4C-E). Given this increase in Tregs, we assessed the apoptotic activity of Treg cells in the spleen but found no significant differences in viability between P2H1^Ad.Cortex^ Treg cells and their WT counterparts (Figure S2G-H). Therefore, this observation suggests that the increase in Tregs may be primarily due to enhanced peripheral differentiation.

### Chronically elevated glucocorticoids reduce B-cell lymphopoiesis

Previously, glucocorticoids have been shown to directly regulate B cell maturation in both the bone marrow and peripheral immune compartments.^38^ In addition, ischemic stroke-induced glucocorticoid signaling has been reported to influence hematopoietic B lineage decisions, leading to impaired adaptive immunity.^39^ Given these established effects, we investigated whether chronic glucocorticoid overproduction in our P2H1^Ad.Cortex^ mice similarly affects B cell development. Consistent with previous findings, our analysis revealed significant disruptions in B cell maturation beyond the pre-pro B cell progenitor stage (lineage^-^ B220^int^ CD93 IgM^-^ CD43 CD24^-^) (Figure 4F). Specifically, we observed a marked reduction in both the number and frequency of pro-B cells (Lin^-^ B220^int^ CD93 IgM^-^ CD19 CD43 CD24^int^), pre-B cells (Lin^-^ B220int CD93 IgM^-^ CD19 CD43^low^ CD24), and immature B cells (Lin^-^ B220^int^ CD93 IgM CD19) in the bone marrow (Figure 4G). These findings suggest that the immunosuppressive effects of chronic levels of glucocorticoids extend to early B cell differentiation and may contribute to the B cell developmental obstruction.

### The GC-GR (GC-receptor) axis regulates myeloid and B-cell differentiation in P2H1^Ad.Cortex^ mice

To directly confirm that the hematopoietic changes observed in P2H1^Ad.Cortex^ mice are driven by glucocorticoid signaling through the GR (GC-GR axis), we performed transplantation experiments using hematopoietic cells deficient in GR (from Vav:cre-GR^f/f^ (GR^HSC^) mice) or WT controls. These cells were transplanted into P2H1^Ad.Cortex^ or littermate controls, as shown in Figure 5A. Twelve months post-transplantation, analysis of B-cell fractions in the spleen revealed that the block in pro-, pre-, and immature B-cell differentiation in P2H1^Ad.Cortex^ mice (Figure 4G) was entirely dependent on GR expression in hematopoietic cells (Figure 5B); an effect that was absent in their transplanted littermate controls (Figure 5C). Similarly, P2H1^Ad.Cortex^ mice receiving GR-deficient BM exhibited a significant reduction in monocytes and PMNs, strongly suggesting that myeloid expansion in this model is also directly regulated by GC-GR signaling (Figure 5D, E). These findings further underscore the significant impact that chronic glucocorticoid overproduction due to HIF1α loss in the adrenal cortex can have on the maturation process of B lymphocytes and myeloid cells.

**Figure 5.**
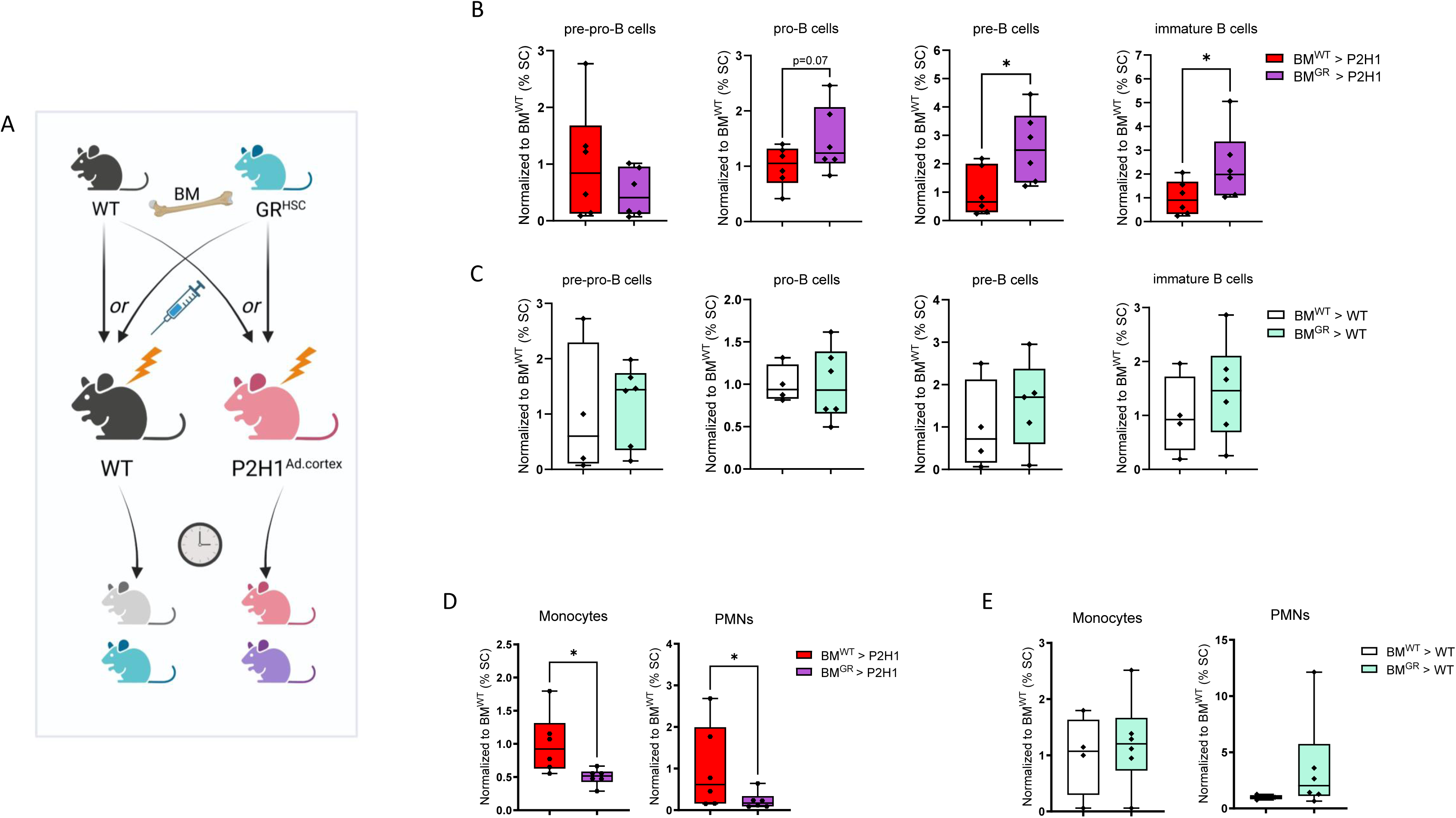
The GC-GR axis regulates myeloid and B-cell differentiation in P2H1^Ad.Cortex^ mice. (A) Schematic overview of the GR^HSC^ or WT BM-transplantation in irradiated P2H1^Ad.Cortex^ or WT recipients. (B-C) Subsequent FACS analysis of different B-cell progenitors in the spleen 12 months after transplantation. (D-E) FACS analysis of monocytes and PMNs. Data points represent individual mice from two different experiments - normalized values against WT control. Data are presented as box & whisker plots showing all data points with whiskers from min to max. Statistical significance was determined using a Mann–Whitney U-test or unpaired t-test with Welch’s correction (*p<0.05).

### HIF1**α**-induced downregulation of steroidogenesis affects erythropoiesis but not HSPCs

In our previous study, we demonstrated that simultaneous deficiency of PHD2 and PHD3 in the adrenal cortex leads to increased HIF1α activity, resulting in a sustained reduction in steroidogenesis, including lower glucocorticoid levels in both the adrenal gland and the circulation.^14^ To further investigate the hematopoietic effects of this condition, we analyzed hematopoietic cells in the P2P3^Ad.Cortex^ mouse line (P2P3), using the same approach as for P2H1^Ad.Cortex^ mice. While elevated glucocorticoids significantly increased HSPC fractions in P2H1^Ad.Cortex^ mice (Figure 1B), we found no significant changes in hematopoietic stem and progenitor cells (HSPCs) in P2P3^Ad.Cortex^ mice compared to WT littermates (Figure 6A and Figure S3A). However, further analysis revealed an increase in RBC levels, accompanied by higher hematocrit (HCT) and hemoglobin (HGB) levels, but lower mean corpuscular volume (MCV) and platelet counts (Figure 6B). Consistent with these findings, secondary erythropoiesis was enhanced in the spleen rather than the bone marrow, correlating with increased spleen weight (Figure 6C and Figure S3B). Analysis of white blood cells (WBCs) and their precursors showed a significant reduction in circulating monocytes, whereas other myeloid cells, T cells, and B cells remained unchanged (Figure 6D and Figure S3C, D). Taken together, these results indicate that P2P3^Ad.Cortex^ mice, despite chronically producing lower levels of glucocorticoids, do not exhibit an inverse phenotype of P2H1^Ad.Cortex^ mice. This underscores the complex interplay between reduced glucocorticoid levels and hematopoiesis, emphasizing that changes in steroidogenesis do not necessarily translate into direct or linear effects on HSPCs and lineage differentiation.

**Figure 6.**
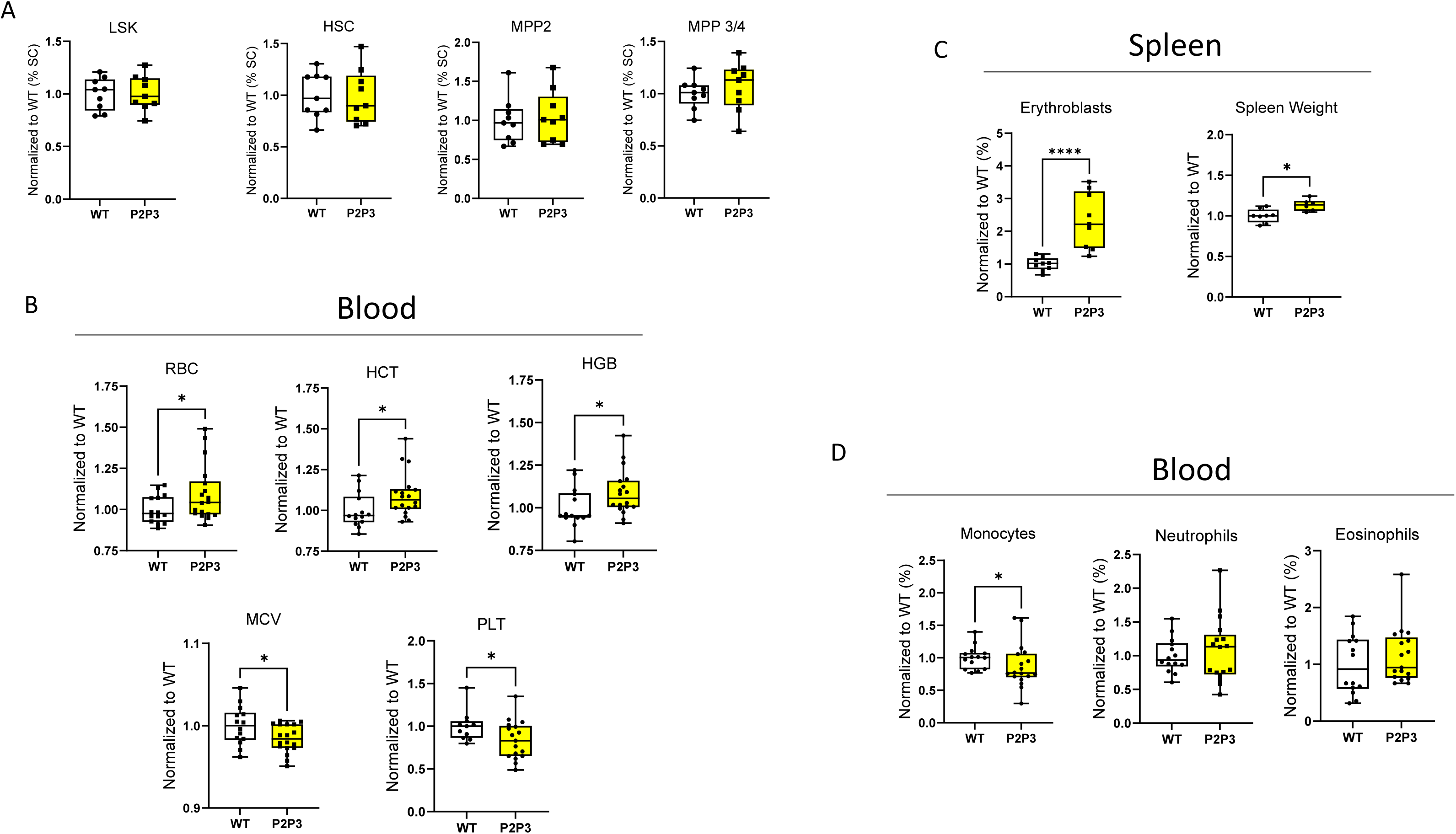
HIF1α-induced downregulation of steroidogenesis affects erythropoiesis but not HSPCs. (A) Normalized percentage of single cells in the BM of WT mice and Akr1b7:cre-PHD2/PHD3^ff/ff^ (P2P3) littermates. Data points represent individual mice from at least three different experiments - normalized values against WT control. Data are represented as mean ± SEM. (B) Normalized number of different RBC parameters in circulation from WT mice and P2P3 littermates under steady state. (C) Representative FACS gating strategy for the identification of erythroblasts and weight of the representative spleen. (D) Sysmex analysis of myeloid cells in the blood (Normalized to the average of WT controls). Data points represent individual mice from two different experiments - normalized values against WT control. Data are presented as box & whisker plots showing all data points with whiskers from min to max. Statistical significance was determined using a Mann–Whitney U-test or unpaired t-test with Welch’s correction (*p<0.05).

## Discussion

Our study demonstrates that chronic glucocorticoid exposure, driven by HIF1α loss in adrenocortical cells in vivo, expands HSPCs while shifting HSCs toward a more quiescent and metabolically restrained state. Functionally, these HSCs exhibited enhanced regenerative potential, while myeloid differentiation was significantly altered, with an increase in GMPs, monocytes and PMNs, but a dramatic inhibition of B cell differentiation. In a transplantation assay using GR-deficient bone marrow into irradiated P2H1^Ad.Cortex^ mice we confirmed that the myeloid and B cell differentiation phenotypes were driven by glucocorticoid receptor (GC-GR) signaling. In contrast, HIF1α-induced downregulation of steroidogenesis in P2P3^Ad.Cortex^ mice had no significant effect on HSPC numbers, B cell differentiation or Treg profiles. Despite lower systemic glucocorticoids, these mice did not exhibit the inverse phenotype of P2H1^Ad.Cortex^ mice, highlighting the complex interplay between steroid hormones and hematopoiesis.

Previous studies on the role of glucocorticoids in HSCs primarily focused on short-term dexamethasone treatment, a synthetic glucocorticoid, which differs from our chronic endogenous approach.^16, 40, 41^ However, consistent with previous experiments using only low doses of dexamethasone ^17^ our analysis revealed increased HSC quiescence, leading to a significant delay in HSC exhaustion, as observed in our long term transplantation experiment. At the molecular level, this prolonged quiescent state appears to be influenced by key regulators that we identified through our RNAseq analysis, which suggest a coordinated network reinforcing chronically mild cell cycle arrest and niche interactions to preserve HSC function under chronic glucocorticoid exposure (p53 pathway activity). Upregulation of *Trp53bp2* likely enhances HSC quiescence through complementary mechanisms. As an enhancer of p53 signaling, Trp53bp2 promotes cell cycle arrest and maintains genomic stability,^42^ potentially preventing HSCs from entering an active proliferative state. This is consistent with the broader role of p53 in enforcing quiescence under conditions of cellular stress including in HSCs.^43^ In parallel, Slamf1 (CD150) is a critical marker of quiescent HSCs, particularly in the long-term stem cell compartment. It is involved in maintaining HSC quiescence by mediating interactions between HSCs and their niche, promoting signals that support dormancy, and they may therefore survive better after transplantation.^44^ The downregulation of *Hsd11b1*, *Mastl*, *E2f1*, and *Cdk5rap1* in HSCs adds up to the coordinated shift toward quiescence. Mastl and E2f1 are both key regulators of the cell cycle, with their suppression leading to inhibition of proliferation—Mastl downregulation increases PP2A-B55, a phosphatase complex that dephosphorylates mitotic substrates, thereby delaying mitotic entry,^45^ while E2f1 suppression reduces the expression of genes required for G1/S transition.^46^ In parallel, Cdk5rap1 downregulation limits mitochondrial activity, promoting a low-energy state that supports dormancy.^47^ Uniquely, Hsd11b1 downregulation in our model may result from chronically elevated systemic glucocorticoid levels, serving as a protective mechanism to limit intracellular glucocorticoid activation and potentially preventing HSC activation.^48^ Together, these molecular changes likely work synergistically to stabilize HSC quiescence in response to chronic glucocorticoid exposure and preserve the long-term integrity of the stem cell pool.

Our P2H1^Ad.Cortex^ mice exhibited marked alterations in mature hematopoietic cells, particularly enhanced erythropoiesis. This mirrors observations in patients with Cushing’s disease, where chronic glucocorticoid exposure is linked to erythrocytosis, elevated hematocrit, and increased hemoglobin levels. Consistent with this, our analysis revealed a significant increase in RBC numbers, hemoglobin levels, and MCV in P2H1^Ad.Cortex^ mice, despite reduced EPO levels. These findings suggest that glucocorticoids directly promote erythropoiesis, operating through an EPO-independent mechanism. This is supported by previous studies indicating that glucocorticoids can directly enhance stress erythropoiesis particularly in the spleen, which aligns with our observation of increased splenic erythroblast populations.^34, 49^ Interestingly, our P2P3^Ad.Cortex^ mice, characterized by downregulated glucocorticoid production, also exhibited a significant increase in RBC counts, although the MCV was significantly reduced in P2P3^Ad.Cortex^ mice compared to wild-type controls. This observation contrasts with typical clinical findings, as conditions of glucocorticoid deficiency, such as in Addison’s disease, are generally associated with anemia rather than erythrocytosis.^50^ Therefore, the mechanisms underlying the increased RBC production in our P2P3^Ad.Cortex^ mice remain unclear and warrant further investigation to elucidate the background of this observed phenotype.

We also observed alterations in myeloid cell populations within the bone marrow and, to a lesser extent, in the spleen. Conversely, eosinophil counts were significantly reduced in these compartments. These findings align with clinical observations in Cushing’s disease, where chronic glucocorticoid excess is associated with elevated myeloid cells, while eosinophil levels are typically decreased, due to suppression of their production.^34^ Furthermore, we observed significant alterations in lymphocyte populations, most notably a severe impairment in B cell development within the bone marrow. Specifically, pro-B, pre-B, and immature B cell populations were significantly reduced, indicating a blockade in B cell maturation beyond the pre-pro-B cell stage. This finding is consistent with reports that glucocorticoids suppress B lymphopoiesis, either through direct effects on B cell progenitors or by modifying the bone marrow microenvironment.^39, 51^

Finally, our transplantation experiments provide decisive evidence that the observed increase in monocytes and PMNs, and the inhibition in B cell maturation are directly mediated by glucocorticoid receptor signaling. The failure of B cell differentiation beyond the pro-pre-B cell stage was entirely dependent on the presence of GR in hematopoietic cells, as this phenotype was absent when GR-deficient HSCs were transplanted in P2H1^Ad.Cortex^ mice. Similarly, mice receiving GR-deficient BM exhibited significantly lower monocyte and PMN numbers, confirming that the myeloid expansion observed in P2H1^Ad.Cortex^ mice is also related to the GR. These findings establish a clear GC-GR-dependent axis regulating both myeloid and B cell lineage commitment, further underscoring the profound impact of low-dose, chronic glucocorticoid overproduction on the hematopoietic system.

Taken together, our findings reveal that chronic glucocorticoid overproduction profoundly alters hematopoiesis, promoting HSPC expansion, increased quiescence, and enhanced regenerative capacity of HSCs, while also shaping lineage differentiation. We demonstrate that GC-GR signaling is a key regulator of myelopoiesis and B lymphopoiesis, driving an increase in monocytes and PMNs, while blocking B cell maturation in the bone marrow. Decisively, our transplantation experiments confirm that these effects are intrinsic to hematopoietic cells and directly mediated by GR signaling. Therefore, this adrenocortical specific HIF1α-deficient mouse model provides a powerful tool for further investigating how chronic glucocorticoid exposure impacts HSC function, immune cell differentiation, and stress hematopoiesis. By dissecting the precise molecular mechanisms underlying these changes, this system may offer valuable insights into glucocorticoid-related hematological disorders and immunosuppressive therapies, ultimately contributing to a better understanding of how HIF-dependent steroid hormone imbalances influence hematopoiesis in health and disease.

## Contributions

D.W., N.E. and M.T.J designed and performed the majority of experiments, analyzed data, and contributed to the writing of the manuscript. T.G. performed experiments, analyzed data and contributed to the discussion/interpretation. C.E., J.T, U.B., M.R., T.C., P.M. and A.E.A. provided tools and contributed to the discussion/interpretation. B.W. designed and supervised the overall study, analyzed data, and wrote the manuscript. The cartoons are created with Biorender.com

## Data-sharing statement

Data is provided within the manuscript or supplementary information files. Materials are available upon reasonable request

## References

1. Wielockx B, Grinenko T, Mirtschink P, Chavakis T. Hypoxia Pathway Proteins in Normal and Malignant Hematopoiesis. Cells. 2019;8(2):

2. Dausinas Ni P, Basile C, Junge C, Hartman M, O’Leary HA. Hypoxia and Hematopoiesis. Current Stem Cell Reports. 2022;8(1):24–34.

3. Simsek T, Kocabas F, Zheng J, et al. The distinct metabolic profile of hematopoietic stem cells reflects their location in a hypoxic niche. Cell Stem Cell. 2010;7(3):380–390.

4. Pollard PJ, Kranc KR. Hypoxia signaling in hematopoietic stem cells: a double-edged sword. Cell Stem Cell. 2010;7(3):276–278.

5. Ding L, Saunders TL, Enikolopov G, Morrison SJ. Endothelial and perivascular cells maintain haematopoietic stem cells. Nature. 2012;481(7382):457–462.

6. Chow A, Lucas D, Hidalgo A, et al. Bone marrow CD169+ macrophages promote the retention of hematopoietic stem and progenitor cells in the mesenchymal stem cell niche. J Exp Med. 2011;208(2):261–271.

7. Zhao JL, Baltimore D. Regulation of stress-induced hematopoiesis. Curr Opin Hematol. 2015;22(4):286–292.

8. Smith MA, Culver-Cochran AE, Adelman ER, et al. TNFAIP3 Plays a Role in Aging of the Hematopoietic System. Front Immunol. 2020;11(536442.

9. Nagai Y, Garrett KP, Ohta S, et al. Toll-like receptors on hematopoietic progenitor cells stimulate innate immune system replenishment. Immunity. 2006;24(6):801–812.

10. Seyfried AN, Maloney JM, MacNamara KC. Macrophages Orchestrate Hematopoietic Programs and Regulate HSC Function During Inflammatory Stress. Front Immunol. 2020;11(1499.

11. Galon J, Franchimont D, Hiroi N, et al. Gene profiling reveals unknown enhancing and suppressive actions of glucocorticoids on immune cells. Faseb j. 2002;16(1):61–71.

12. Cain DW, Cidlowski JA. Immune regulation by glucocorticoids. Nature Reviews Immunology. 2017;17(4):233–247.

13. Rhen T, Cidlowski JA. Antiinflammatory action of glucocorticoids--new mechanisms for old drugs. N Engl J Med. 2005;353(16):1711–1723.

14. Watts D, Stein J, Meneses A, et al. HIF1α is a direct regulator of steroidogenesis in the adrenal gland. Cell Mol Life Sci. 2021;78(7):3577–3590.

15. Rawat S, Dadhwal V, Mohanty S. Dexamethasone priming enhances stemness and immunomodulatory property of tissue-specific human mesenchymal stem cells. BMC Dev Biol. 2021;21(1):16.

16. Kim H, Choi JY, Lee JM, et al. Dexamethasone increases angiopoietin-1 and quiescent hematopoietic stem cells: a novel mechanism of dexamethasone-induced hematoprotection. FEBS Lett. 2008;582(23-24):3509–3514.

17. Eka Putra W, Soewondo A, Rifa’i M. Effect of Dexamethasone Administration toward Hematopoietic Stem Cells and Blood Progenitor Cells Expression on BALB/c Mice. The Journal of Pure and Applied Chemistry Research. 2015;4(3):100–108.

18. Byron JW. Effect of Steroids on the Cycling of Haemopoietic Stem Cells. Nature. 1970;228(5277):1204–1204.

19. Semenza GL. Pharmacologic Targeting of Hypoxia-Inducible Factors. Annu Rev Pharmacol Toxicol. 2019;59(379–403.

20. Mittelstadt PR, Monteiro JP, Ashwell JD. Thymocyte responsiveness to endogenous glucocorticoids is required for immunological fitness. J Clin Invest. 2012;122(7):2384–2394.

21. Stadtfeld M, Graf T. Assessing the role of hematopoietic plasticity for endothelial and hepatocyte development by non-invasive lineage tracing. Development. 2005;132(1):203–213.

22. Singh RP, Grinenko T, Ramasz B, et al. Hematopoietic Stem Cells but Not Multipotent Progenitors Drive Erythropoiesis during Chronic Erythroid Stress in EPO Transgenic Mice. Stem cell reports. 2018;10(6):1908–1919.

23. Singh RP, Franke K, Kalucka J, et al. HIF prolyl hydroxylase 2 (PHD2) is a critical regulator of hematopoietic stem cell maintenance during steady-state and stress. Blood. 2013;121(26):5158–5166.

24. Martin M. Cutadapt removes adapter sequences from high-throughput sequencing reads. 2011. 2011;17(1):3.

25. Dobin A, Davis CA, Schlesinger F, et al. STAR: ultrafast universal RNA-seq aligner. Bioinformatics. 2012;29(1):15–21.

26. Anders S, Huber W. Differential expression analysis for sequence count data. Genome Biology. 2010;11(10):R106.

27. Anders S, Pyl PT, Huber W. HTSeq—a Python framework to work with high-throughput sequencing data. Bioinformatics. 2014;31(2):166–169.

28. Wickham H. ggplot2: elegant graphics for data analysis. New York Springer-Verlag, 2016.

29. Gu Z, Eils R, Schlesner M. Complex heatmaps reveal patterns and correlations in multidimensional genomic data. Bioinformatics. 2016;32(18):2847–2849.

30. Subramanian A, Tamayo P, Mootha VK, et al. Gene set enrichment analysis: a knowledge-based approach for interpreting genome-wide expression profiles. Proc Natl Acad Sci U S A. 2005;102(43):15545–15550.

31. Alhamdoosh M, Ng M, Wilson NJ, et al. Combining multiple tools outperforms individual methods in gene set enrichment analyses. Bioinformatics. 2017;33(3):414–424.

32. Schubert M, Klinger B, Klünemann M, et al. Perturbation-response genes reveal signaling footprints in cancer gene expression. Nature Communications. 2018;9(1):20.

33. Crochemore C, Michaelidis TM, Fischer D, Loeffler JP, Almeida OF. Enhancement of p53 activity and inhibition of neural cell proliferation by glucocorticoid receptor activation. FASEB J. 2002;16(8):761–770.

34. Varricchio L, Geer EB, Martelli F, et al. Patients with hypercortisolemic Cushing disease possess a distinct class of hematopoietic progenitor cells leading to erythrocytosis. Haematologica. 2023;108(4):1053–1067.

35. Franke K, Kalucka J, Mamlouk S, et al. HIF-1alpha is a protective factor in conditional PHD2-deficient mice suffering from severe HIF-2alpha-induced excessive erythropoiesis. Blood. 2013;121(8):1436–1445.

36. Jia WY, Zhang JJ. Effects of glucocorticoids on leukocytes: Genomic and non-genomic mechanisms. World J Clin Cases. 2022;10(21):7187–7194.

37. Cari L, De Rosa F, Nocentini G, Riccardi C. Context-Dependent Effect of Glucocorticoids on the Proliferation, Differentiation, and Apoptosis of Regulatory T Cells: A Review of the Empirical Evidence and Clinical Applications. Int J Mol Sci. 2019;20(5):

38. Gruver-Yates AL, Quinn MA, Cidlowski JA. Analysis of glucocorticoid receptors and their apoptotic response to dexamethasone in male murine B cells during development. Endocrinology. 2014;155(2):463–474.

39. Courties G, Frodermann V, Honold L, et al. Glucocorticoids Regulate Bone Marrow B Lymphopoiesis After Stroke. Circ Res. 2019;124(9):1372–1385.

40. Pandolfi J, Baz P, Fernández P, et al. Regulatory and effector T-cells are differentially modulated by Dexamethasone. Clinical Immunology. 2013;149(3, Part B):400–410.

41. Prado C, Gómez J, López P, de Paz B, Gutiérrez C, Suárez A. Dexamethasone upregulates FOXP3 expression without increasing regulatory activity. Immunobiology. 2011;216(3):386–392.

42. Liu G, Parant JM, Lang G, et al. Chromosome stability, in the absence of apoptosis, is critical for suppression of tumorigenesis in Trp53 mutant mice. Nat Genet. 2004;36(1):63–68.

43. Liu Y, Elf SE, Asai T, et al. The p53 tumor suppressor protein is a critical regulator of hematopoietic stem cell behavior. Cell Cycle. 2009;8(19):3120–3124.

44. Wilson A, Oser GM, Jaworski M, et al. Dormant and self-renewing hematopoietic stem cells and their niches. Ann N Y Acad Sci. 2007;1106(64–75.

45. Wong PY, Ma HT, Lee HJ, Poon RY. MASTL(Greatwall) regulates DNA damage responses by coordinating mitotic entry after checkpoint recovery and APC/C activation. Sci Rep. 2016;6(22230.

46. Ma Y, Croxton R, Moorer RL, Jr., Cress WD. Identification of novel E2F1-regulated genes by microarray. Arch Biochem Biophys. 2002;399(2):212–224.

47. Xie Q, Wu Q, Horbinski CM, et al. Mitochondrial control by DRP1 in brain tumor initiating cells. Nat Neurosci. 2015;18(4):501–510.

48. Harris HJ, Kotelevtsev Y, Mullins JJ, Seckl JR, Holmes MC. Intracellular regeneration of glucocorticoids by 11beta-hydroxysteroid dehydrogenase (11beta-HSD)-1 plays a key role in regulation of the hypothalamic-pituitary-adrenal axis: analysis of 11beta-HSD-1-deficient mice. Endocrinology. 2001;142(1):114–120.

49. Gursoy A, Dogruk Unal A, Ayturk S, et al. Polycythemia as the first manifestation of Cushing’s disease. J Endocrinol Invest. 2006;29(8):742–744.

50. Isidori AM, Minnetti M, Sbardella E, Graziadio C, Grossman AB. Mechanisms in endocrinology: The spectrum of haemostatic abnormalities in glucocorticoid excess and defect. European journal of endocrinology / European Federation of Endocrine Societies. 2015;173(3):R101–113.

51. Igarashi H, Medina KL, Yokota T, et al. Early lymphoid progenitors in mouse and man are highly sensitive to glucocorticoids. Int Immunol. 2005;17(5):501–511.

