## Supplementary Figures + Table for "Hypoxia Inducible Factor 1α-driven steroidogenesis impacts systemic hematopoiesis"

Supplementary Figure 1

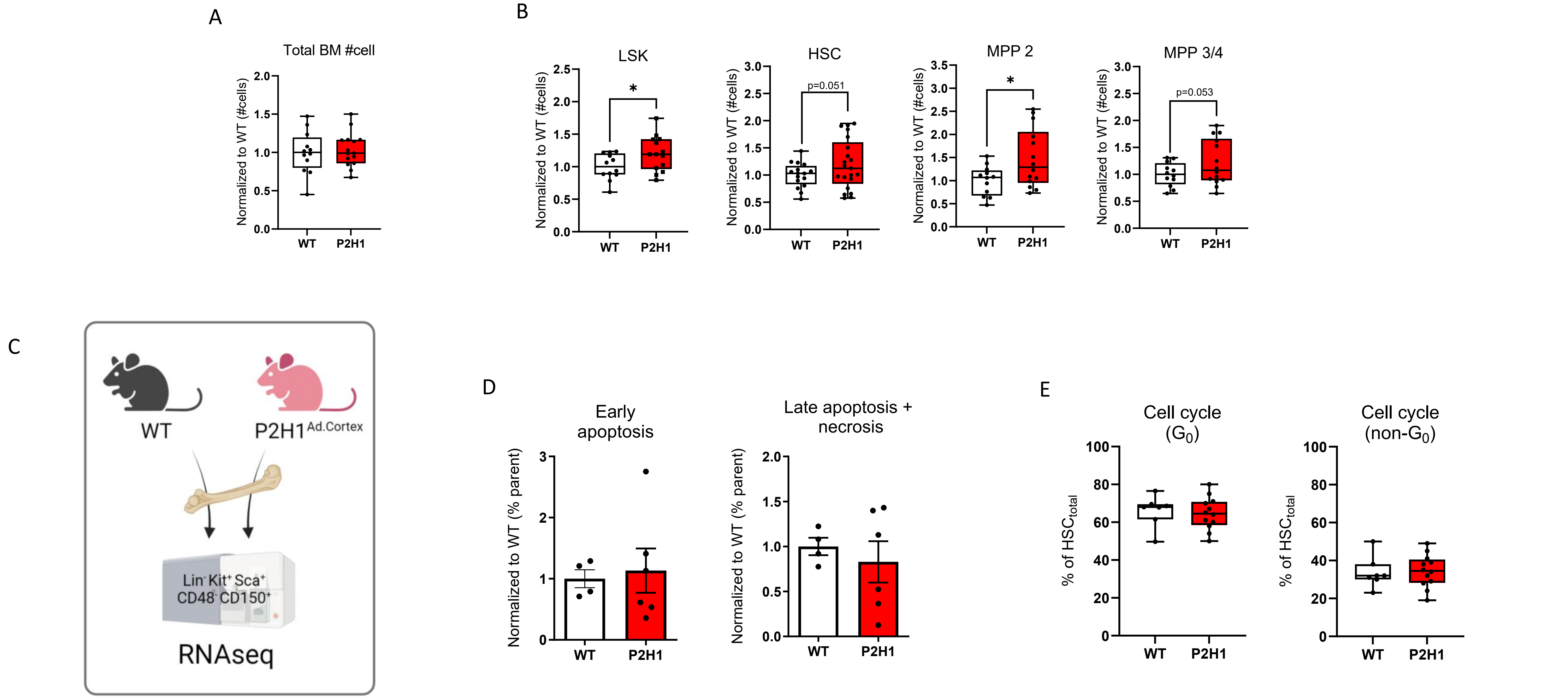


**Supplementary Figure 1. HIF1α-associated steroidogenesis affects the number of hematopoietic stem and progenitor cells.** (A) Total bone marrow cell number. (B) Normalized number of LSK and three different HSPC populations in the bone marrow from WT mice and P2H1 littermates. Each graph represents data from at least three independent experiments. (C) Schematic overview of the RNAseq approach representing the sorting of HSC cells from the bone marrow of P2H1 and WT littermates. (D) Apoptosis (early/late)/Necrosis of HSCs (from 2 independent experiments) and (E) HSC fractions in $G_0$ or non-$G_0$ phases of the cell cycle measured via FACS (from 3 independent experiments). Data are presented either as mean ± SEM or as box & whisker plots showing all data points with whiskers from min to max. Statistical significance was determined using a Mann–Whitney U-test or unpaired t-test with Welch's correction (*p<0.05).

Supplementary Figure 2

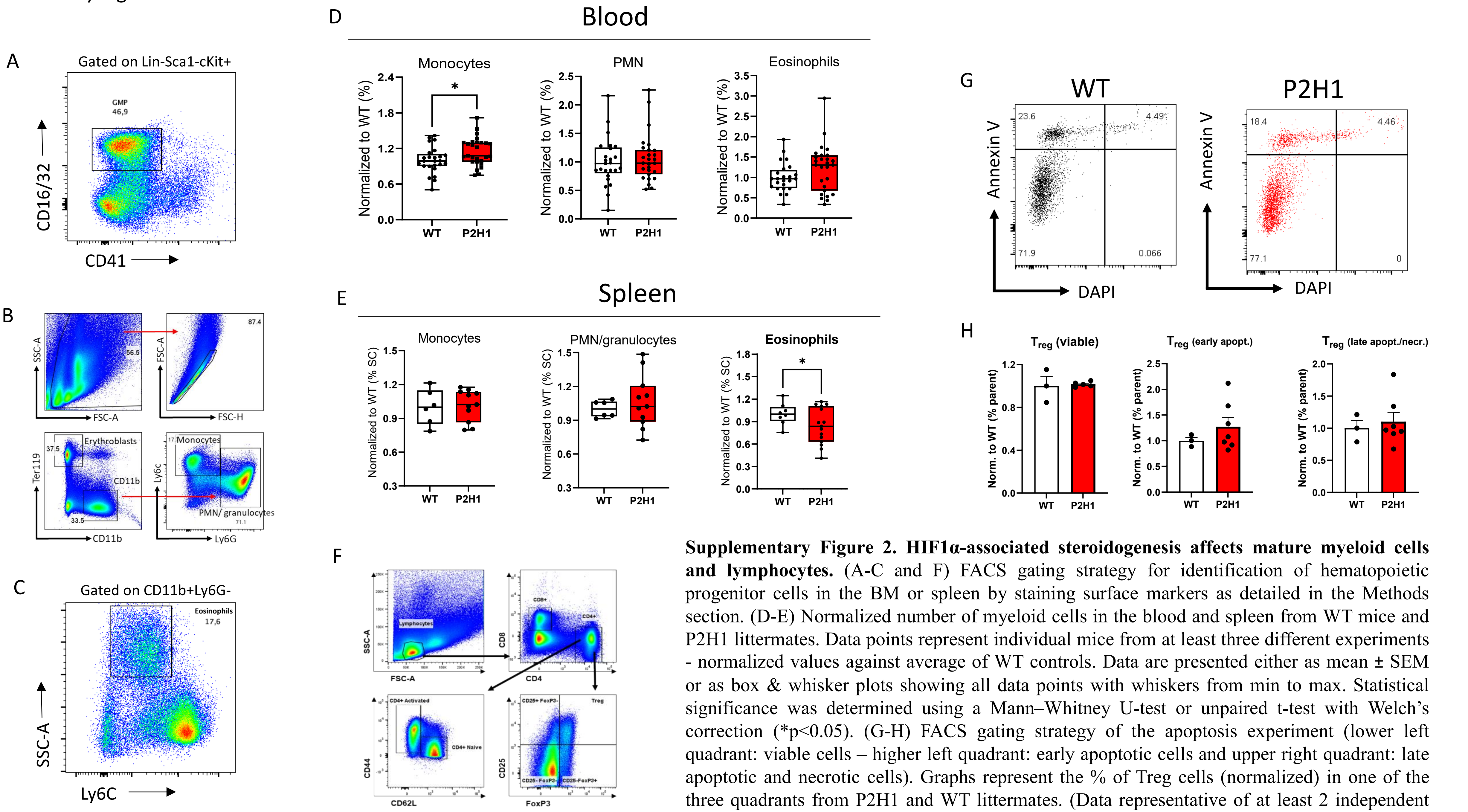


**Supplementary Figure 2. HIF1α-associated steroidogenesis affects mature myeloid cells and lymphocytes.** (A-C and F) FACS gating strategy for identification of hematopoietic progenitor cells in the BM or spleen by staining surface markers as detailed in the Methods section. (D-E) Normalized number of myeloid cells in the blood and spleen from WT mice and P2H1 littermates. Data points represent individual mice from at least three different experiments - normalized values against average of WT controls. Data are presented either as mean ± SEM or as box & whisker plots showing all data points with whiskers from min to max. Statistical significance was determined using a Mann–Whitney U-test or unpaired t-test with Welch's correction (*$p<0.05$). (G-H) FACS gating strategy of the apoptosis experiment (lower left quadrant: viable cells – higher left quadrant: early apoptotic cells and upper right quadrant: late apoptotic and necrotic cells). Graphs represent the % of Treg cells (normalized) in one of the three quadrants from P2H1 and WT littermates. (Data representative of at least 2 independent experiments)

Supplementary Figure 3

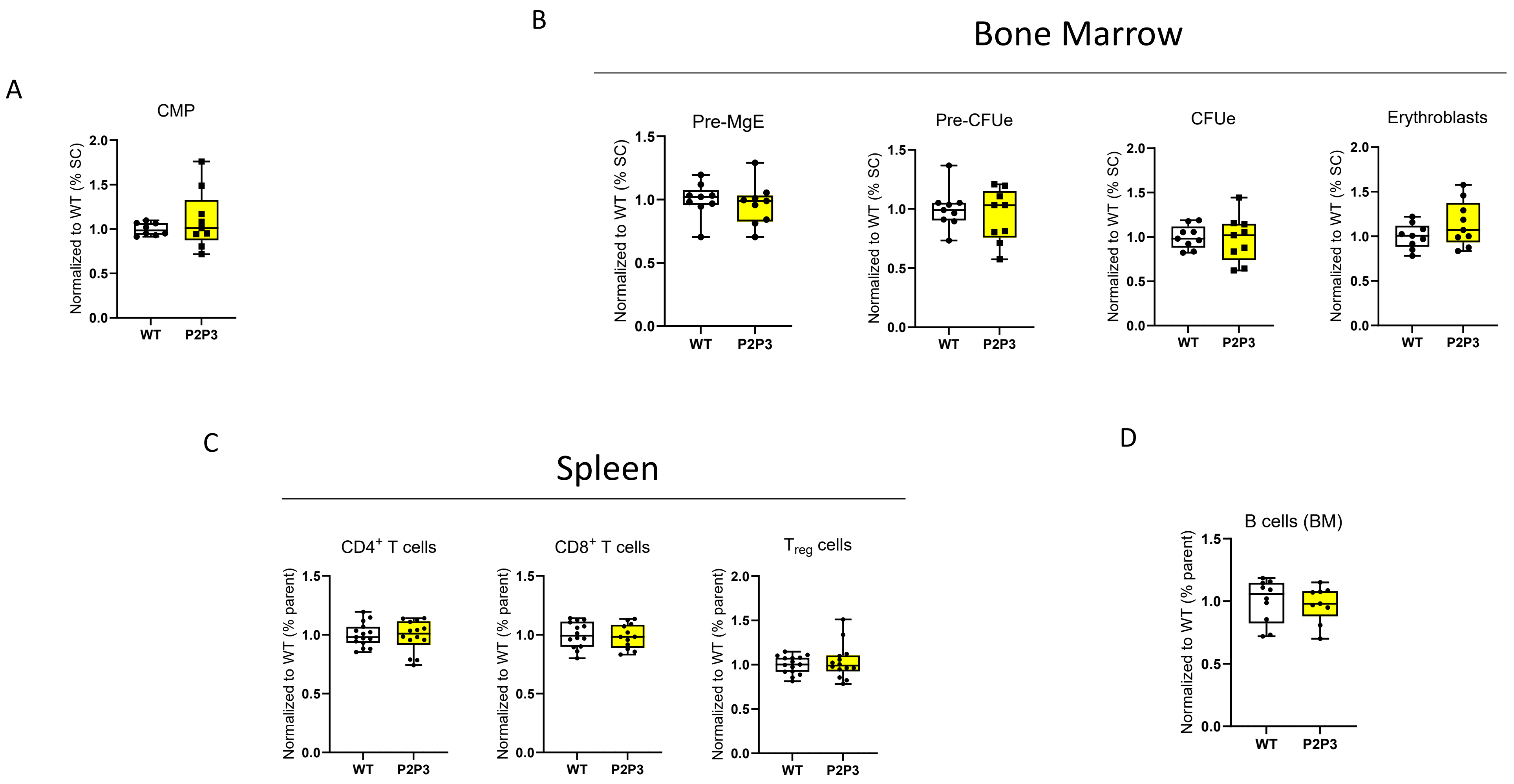


**Supplementary Figure 3. HIF1α reduced steroidogenesis marginally affects the number of hematopoietic progenitor cells and mature cells.** (A-D) FACS analysis of mature hematopoietic populations and progenitors in BM or spleen. Data points represent individual mice from at least two different experiments - normalized values against WT control. Data are presented as box & whisker plots showing all data points with whiskers from min to max.

**Supplementary Table 1: Overview of the antibodies used in the different experiments**

| Antibodies | Host | Cat. Number | Company |
|---|---|---|---|
| CD3e Monoclonal Antibody (145-2C11), Biotin, eBioscience™ (1:1000) | Armenian Hamster | 13-0031-82 | Invitrogen |
| CD19 Monoclonal Antibody (eBio1D3 (1D3)), Biotin, eBioscience™ (1:500) | Rat | 13-0193-81 | Invitrogen |
| NK1.1 Monoclonal Antibody (PK136), Biotin, eBioscience™ (1:2000) | Mouse | 13-5941-81 | Invitrogen |
| TER-119 Monoclonal Antibody (TER-119), Biotin (1:200) | Rat | MA5-17819 | Invitrogen |
| CD11b Monoclonal Antibody (M1/70), Biotin, eBioscience™ (1:500) | Rat | 13-0112-81 | Invitrogen |
| Ly-6G/Ly-6C Monoclonal Antibody (RB6-8C5), Biotin, eBioscience™ (1:800) | Rat | 13-5931-82 | Invitrogen |
| CD45R (B220) Monoclonal Antibody (RA3-6B2), Biotin, eBioscience™ (1:400) | Rat | 13-0452-82 | Invitrogen |
| CD16/CD32 Monoclonal Antibody (93), Alexa Fluor™ 700, eBioscience™ (1:50) | Rat | 56-0161-82 | Invitrogen |
| APC anti-mouse CD48 Antibody (1:300) | Armenian Hamster | 103412 | BioLegend |
| PE/Cyanine7 anti-mouse CD150 (SLAM) Antibody (1:100) | Rat | 115914 | BioLegend |
| CD117 (c-Kit) Monoclonal Antibody (2B8), APC-eFluor™ 780, eBioscience™ (1:600) | Rat | 47-1171-80 | Invitrogen |
| Ly-6A/E (Sca-1) Monoclonal Antibody (D7), PE-Cyanine5, eBioscience™ (1:100) | Rat | 15-5981-82 | Invitrogen |
| CD105 (Endoglin) Monoclonal Antibody (MJ7/18), PE, eBioscience™ (1:400) | Rat | 12-1051-82 | Invitrogen |
| CD34 Monoclonal Antibody (RAM34), FITC, eBioscience™ (1:50) | Rat | 11-0341-85 | Invitrogen |
| CD41a Monoclonal Antibody (eBioMWReg30 (MWReg30)), PerCP-eFluor™ 710, eBioscience™ (1:400) | Rat | 46-0411-82 | Invitrogen |
| eBioscience™ Streptavidin eFluor™ 450 Conjugate (1:300) | | 48-4317-82 | Invitrogen |
| CD3e Monoclonal Antibody (eBio500A2 (500A2)), Alexa Fluor™ 700, eBioscience™ (1:100) | Armenian Hamster | 56-0033-82 | Invitrogen |
| CD45R (B220) Monoclonal Antibody (RA3-6B2), PE-Cyanine7, eBioscience™ (1:500) | Rat | 25-0452-82 | Invitrogen |
| CD11b Monoclonal Antibody (M1/70), eFluor™ 450, eBioscience™ (1:800) | Rat | 48-0112-80 | Invitrogen |
| Ly-6G Monoclonal Antibody (1A8-Ly6g), APC, eBioscience™(1:200) | Rat | 17-9668-80 | Invitrogen |
| BD Pharmingen™ FITC Rat Anti-Mouse Ly-6C (1:300) | Rat | 553104 | BD |
| TER-119 Monoclonal Antibody (TER-119), PE-Cyanine5, eBioscience™ (1:100) | Rat | 15-5921-83 | Invitrogen |
| F4/80 Monoclonal Antibody (BM8), PE, eBioscience™ (1:100) | Rat | 12-4801-82 | Invitrogen |
| CD3 Monoclonal Antibody (17A2), APC, eBioscience™ (1:200) | Armenian Hamster | 17-0032-82 | Invitrogen |

| | | | |
|---|---|---|---|
| CD4 Monoclonal Antibody (GK1.5), PE, eBioscience™ (1:200) | Rat | 12-0041-82 | Invitrogen |
| CD8a Monoclonal Antibody (53-6.7), eFluor™ 506, eBioscience™ (1:400) | Rat | 69-0081-82 | Invitrogen |
| CD25 Monoclonal Antibody (PC61.5), PE-Cyanine7, eBioscience™ (1:100) | Rat | 25-0251-81 | Invitrogen |
| Pacific Blue™ anti-mouse CD62L Antibody (1:200) | Rat | 104423 | BioLegend |
| APC/Cyanine7 anti-mouse/human CD44 Antibody (1:200) | Rat | 103028 | BioLegend |
| FOXP3 Monoclonal Antibody (FJK-16s), FITC, eBioscience™ (1:100) | Rat | 11-5773-82 | Invitrogen |
| CD71 (Transferrin Receptor) Monoclonal Antibody (R17217 (RI7 217.1.4)), FITC, eBioscience™ (1:200) | Rat | 11-0711-81 | Invitrogen |
| Alexa Fluor® 647 anti-mouse TER-119/Erythroid Cells Antibody (1:200) | Rat | 116218 | Biolegend |
| CD45R (B220) Monoclonal Antibody (RA3-6B2), Alexa Fluor™ 700, eBioscience™ (1:100) | Rat | 56-0452-82 | Invitrogen |
| CD93 (AA4.1) Monoclonal Antibody (AA4.1), PE, eBioscience™ | Rat | 12-5892-82 | Invitrogen |
| APC anti-mouse IgM Antibody (1:100) | Rat | 406509 | BioLegend |
| CD19 Monoclonal Antibody (eBio1D3 (1D3)), eFluor™ 506, eBioscience™ (1:100) | Rat | 69-0193-82 | Invitrogen |
| PE/Cyanine7 anti-mouse CD43 Antibody (1:400) | Rat | 143210 | BioLegend |
| CD24 Monoclonal Antibody (M1/69), FITC, eBioscience™ | Rat | 11-0242-82 | Invitrogen |
